# Post-transcriptional dysregulation in autism, schizophrenia and bipolar disorder

**DOI:** 10.1101/2021.02.28.433176

**Authors:** Yuanyuan Wang, Liya Liu, Mingyan Lin

## Abstract

Post-transcriptional gene regulation (PTGR) contributes to numerous aspects of RNA metabolism. While multiple regulators of PTGR have been associated with the occurrence and development of psychiatric disorders, a systematic investigation of the role of PTGR in the context of neuropsychiatric disorders is still lacking. In this work, we developed a new transcriptome -based algorithm to estimate PTGR and applied it to an RNA-Seq dataset of 2160 brain samples from individuals with autism spectrum disorder (ASD), schizophrenia (SCZ), bipolar disorder (BD) and controls. The results showed that the contribution of PTGR abnormality to gene differential expression between three common psychiatric disorders and controls was about 30% of that of transcriptional gene regulation (TGR) abnormality. Besides, aberrant PTGR tended to decrease RNA stability in SCZ/BD, while increase RNA stability in ASD, implicating contrasting pathologies among diseases. The abnormal alteration of PTGR in SCZ/BD converged on the inhibition of neurogenesis and neural differentiation, whereas dysregulation of PTGR in ASD induced enhanced activity of apoptosis. This suggested that heterogeneity in disease mechanism and clinical manifestation across different psychiatric disorders may be partially attributed to the diverse role of PTGR. Intriguingly, we identified a promising RBP (RNA bind protein) ELAVL3 (ELAV-Like Protein 3) that have a profound role in all three psychiatric disorders. Our systematic study expands the understanding of the link between PTGR and psychiatric disorders and also open a new avenue for deciphering the pathogenesis of psychiatric disorders.

## Introduction

Psychiatric disorders, characterized by brain dysfunction, are caused by a combination of psychological, biological, and environmental factors, including schizophrenia (SCZ), bipolar disorder (BD), depression (Depression) and autism (ASD), and so on. Currently, it is generally postulated that psychiatric disorders are polygenic genetic disease with high genetic heterogeneity and complexity, the pathological mechanisms of psychiatric disorders remain poorly understood^1^. In the present study, several studies have revealed that the majority of the psychiatric associated risk loci lies in the gene regulatory region (non-coding region of gene) rather than loci within the gene coding region^2-4^, indicating that a greater role for dysregulation of gene expression and alternative splicing compared to dysfunction of the gene in contributing to the development of psychiatric disorders. Gene expression and alternative splicing are jointly determined by the transcriptional regulation mechanism (TGR) and the post-transcriptional regulation mechanism (PTGR). Recognizing the importance of understanding gene expression and splicing, several researches have undertaken large-scale efforts to study the impact of TGR abnormalities on psychiatric disorders^5,6^, and most of these studies focused on dysregulation of transcription factors and epigenetic patterns^7,8^. These investigations reported several susceptibility factors including the SCZ high risk *ZNF804A*^9^, 22q11.2 microdeletion^10^, ASD high risk *CHD*8^11^, as well as the fever in maternal immune activation^12^. Meanwhile, PTGR, especially RBPs-mediated PTGR, as a major contributor in the control of gene expression and splicing regulation, and participates in the regulation of RNA metabolism^13,14^. Nevertheless, there are few studies on the PTGR estimation algorithm, leaving large parts of the PTGR, especially RBPs-mediated PTGR, unexplored^15^. A more comprehensive analysis focusing on the PTGR and psychiatric disorders is necessary to accelerate our understanding of the role of RBPs-mediated PTGR in the pathogenesis of psychiatric disorders.

Several studies based on RNA-seq provided evidence that the reads in intronic regions can represent the expression level of pre-mRNAs, which are mainly transcriptionally driven^16-18^. The reads in the exonic regions are correlated with the expression level of mature mRNAs (mature mRNAs), which are related to transcriptional in combination with post-transcriptional regulation events^16-18^. Therefore, the changes of the expression level of pre-mRNAs and mature mRNAs across different conditions allow us to evaluate the contributions of post-transcriptional regulation to observed changes in steady-state RNA levels and identify abnormal changes of gene post-transcriptional regulatory mechanism. Thus, these analytic methods have defined the difference of the logarithm of fold-change of exonic reads and the logarithm of fold-change of intronic reads (Δexon–Δintron) as altered post-transcriptional regulation of mRNAs^16-18^. It is known that Δexon and Δintron are also affected by the RNA metabolic limitations, measurement errors, and other factors, and the presence of these biases leads to ∆intron much larger than ∆exon. However, these methods do not take into account these biases, which results in the overestimation of ∆PTGR.

To provide a comprehensive view of the abnormal PTGR architecture across three common psychiatric disorders (ASD, SCZ, and BD) at the whole-genome level and measure the contributions of RBPs-mediated PTGR for disease-specific changes in gene expression, we developed a computational method systematically analyzed the PTGR changes of 2160 brain transcriptome samples data from the PsychENCODE Consortium^19^, including 80 ASD samples, 594 SCZ samples, and 253 BD samples, as well as control. In summary, our analyses offer an updated method for PTGR estimation, and highlight that abnormal PTGR as a potential mechanism conferring key aspects of psychiatric disorders specificity and complexity, and find a novel direction for further research into the etiology of psychiatric disorders.

## Materials and Methods

### Data collection

For this study, all processed bulk RNA-seq alignment (.bam) files were collected from the PsychENCODE consortium. All bam files were aligned to the hg19 reference genome and available at https://www.synapse.org/#!Synapse:syn6039873. These include normal controls (n=1233), as well as schizophrenia (SCZ) (n=594), bipolar disorder (BD) (n=253), autism spectrum disorder (ASD) (n=80).

### Annotation of exon and intron reads

For reads annotation, we made a custom intron GTF annotation file and a custom composite exon GTF annotation file based on gencode v19 GTF. The following set of rules was applied to generate two custom GTF files supporting a given genomic region as an exonic region or intronic region: intron coordinates of genes were extracted from GENCODE v19 as custom intronic GTF; exon and untranslated region coordinates within any isoform of genes were extracted from GENCODE v19 as custom composite exonic GTF. Gene-level read counts were calculated for intron and exon separately using featureCounts^20^.

### Covariate selection

Exon/intron counts were compiled from gene level imported into R for downstream analyses. Genes were quantified in TPM and further filtered to include those with TPM > 1 in at least 50% of samples.

Bulk RNA-Seq samples clinical data are available at https://www.synapse.org/#!Synapse:syn4587614. Missing values in clinical data were imputed using the *missMDA* R package^21^. We summarized the top 40 principal components which collectively explained 99% of the total variance. To determine which covariates to include in the final differential post-transcriptional regulation model, we performed multivariate adaptive regression as implemented in the *earth* package in R for intronic and exonic separately. The superset of potential covariates available for all samples included: study, tissue, libraryPrep, strand specificity, platform, individual ID Source, diagnosis, sex, ethnicity, PMI, RIN, ageDeath, along with all 40 seqPCs. For continuous variables, we also included squared terms. These covariates with input into the *earth* model along with count data (limma voom normalized, centered, and scaled). The model was run using linear predictors and otherwise default parameters. As the model fits a maximum of 1000 genes simultaneously, we performed 1000 permutations randomly subsetting 1000 genes at a time. From this, we chose as a set of known covariates those present in a total of the exons and introns resulting pruned models, which consisted of: study, diagnosis, sex, ethnicity, PMI, ageDeath, PC (1-3), PC (5-11), PC (13-18), PC20, PC22, PC23, PC (25-27), PC29, PC31, PC (33-36), PC39, RINS2, ageDeathS2, PC1S2, PC2S2, PC4S2, PC6S2, PC9S2, PC16S2, PC17S2, PC22S2.

### Exonic/Intronic count correction

Count level quantifications were normalized for library size using TMM normalization in *edgeR* and were transformed as log2(CPM+0.5). Normalized and transformed exon/intron counts were then calculated using a linear mixed-effects model using the *nlme* package in R. The covariates specified in the previous section were included as fixed effects in the model. In addition, we included a random effect term for each unique subject to account for subject overlap across sequencing studies. All covariates except for diagnosis and subject were regressed from our exonic/intronic count dataset.

### New PTGR estimation algorithm

The intronic counts representing the relative abundance of pre-mRNA, and the exonic counts representing the relative abundance of mature-mRNA. We took the median of all samples within covariates corrected exonic/intronic counts dataset as the reference sample *S*ref, and then for each gene in each sample *S*, we subtracted the median value across the samples to obtain Δexon, Δintron. Different condition exonic and intronic groups were analyzed separately.

For each of control, SCZ, ASD, and BD set separately, we modeled the ratio of Δexon and Δintron for each gene by robust linear regression of Δexon vs. Δintron. The control group ratio is recorded as slope A. We used the same method to estimate the ratio of ∆exon and ∆intron in the disease group. The disease group ratio is recorded as slope B. If B>A, which means that PTGR is up-regulated in the B(disease) group (∆PTGR > 0), and representing the increased RNA stability in the disease group. Positive and negative ∆PTGR values correspond to increased RNA stability and reduced RNA stability, respectively. Statistical significance of ∆PTGR was determined using *t-test* shown below:

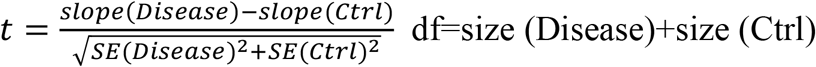

Here, *Slope* represents the robust linear regression slope value, and *SE* represents standard error of *Slope*. The df is the sum of samples in the control and disease groups.

For each gene, the effect of PTGR vs TGR (transcription gene regulation) on gene expression can be estimated according to the following formula:

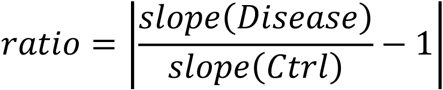

The average ∆PTGR of each gene was obtained from the above analysis. To measure the ∆PTGR (∆ PTGR_in) of each gene in each individual of the disease, we used the following formula:

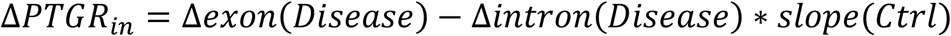

### Post-transcriptional perturbation dataset analysis

We used RNA-seq data of schizophrenia patient-derived neural progenitor cells with purely post-transcriptional perturbations mediated by microRNAs from ref^22^. (GEO accession GSE80170) to evaluate the performance of the PTGR algorithm. The data include control samples, samples with the miR-9 knockdown, and samples with the miR-9 overexpression. Reads were mapped to the hg19 assembly of human genome using STAR^23^ with default parameters. We used custom intronic GTF and custom composite exonic GTF for reads annotation, and gene-level read counts were calculated for intronic and exonic separately using featureCounts. We identified genes that were inferred to be up-PTGR or down-PTGR in the knockdown/overexpression group. Because direct targets of miRNAs are enriched in miRNA seed matches in their 3′-UTR, we examined whether is genes that were up-PTGR or down-PTGR were enriched for miR-9 RNA seed sites in their 3′UTRs to evaluate the performance of PTGR algorithm.

### REMBRA PTGR effect deconvolution algorithm

We closely followed the REMBRA pipeline for PTGR effect deconvolution. We did this to optimally enable comparisons between our PTGR algorithm and REMBRA algorithm. These steps are implemented in a software package available at https://github.com/csglab/REMBRANDTS^17^.

### Integrated PsychENCODE transcription regulatory results

Summary differential expression genes results from the PsychENCODE were obtained from http://resource.psychencode.org/DER-13_Disorder_DEX_Genes.csv.

### Identify key RBPs

We predicted the interaction between DPRGs and 195 human RBPs through the combination of RBPs and phantoms, and then inferred the RBPs that plays a key role through linear models. We limited the analysis of RBP and miRNA binding sites to genes for which all isoforms had the same 3′ UTR start coordinates, and the 3’ UTR was composed of a single exon, in order to minimize the possibility of confounding effects of alternative splicing. The 3′ UTR start coordinates were extracted from GENCODE v19. For analysis of RBP binding sites, we first collected a non-redundant compendium of available sequence preferences for human RBPs. We obtained 217 position frequency matrices (PFMs) representing 117 human RBPs from beRBP^24^, including PFMs with direct experimental evidence and those inferred by homology. Then, we calculated all pairwise PFM similarities using MoSBAT^25^, and then used affinity propagation to cluster the PFMs based on similarity, keeping only the “exemplar” from each cluster. This reduced the total number of PFMs to 154, which we call the non-redundant RBP motif set. Then, we scanned the 3′ UTR sequences with each PFM using AffiMx from the MoSBAT package, resulting in a vector of PFM scores that represents the affinity of the corresponding RBP for binding to different 3′ UTRs.

To identify RBPs that are associated with disease-specific stability profiles, we used the gene-level stability measures as the response variable in multiple linear regression, with RBP binding site match counts as predictor variables. RBPs and miRNAs whose binding sites were significantly associated with disease-specific stability were identified based on t-test of regression coefficients at *pvalue* < 0.05.

### Developmental expression data

Developmental expression data from the BrainSpan: Atlas of the Developing Human Brain (http://www.brainspan.org/).

### Functional enrichment analysis

We performed gene ontology (GO) enrichment analysis using the R package *clusterProfiler*^26^ and GO interactome network was performed by Metascape^27^.

## Results

### New PTGR evaluation algorithm

For an unbiased estimate of the post-transcriptional regulation contributions to gene expression changes, we devised a computational method and applied it to the standard RNA-seq experiment. The abundance of reads lies in the intron region in RNA-seq data correspond to pre-mature mRNA abundance, changes in the abundance of intronic reads can be used to estimate the change in transcription regulation. While the abundance of reads lies in the exon region in RNA-seq data correspond to mature mRNA abundance, changes in the abundance of exonic reads can be used to estimate the changes in transcription regulation and post-transcription regulation. (Methods and Fig. 1A). We first quantified the abundance of exonic and intronic reads in PsychENCODE RNA-seq datasets. As expected, we found that about 38% of reads were intronic reads, and 62% of reads were exonic reads (Fig. 1A). This observation is in agreement with previous studies^16,18^, a substantial number of reads in bulk RNA-Seq originate from pre-mature mRNA. For each of control, SCZ, ASD, and BD set separately, we modeled the ratio of Δexon and Δintron for each gene by robust linear regression of Δexon vs Δintron (Methods and Fig. 1B). The control group ratio is recorded as slope A. We used the same method to estimate the ratio of ∆exon and ∆intron in the disease group. The disease group ratio is recorded as slope B. If B>A, which means that the PTGR is up-regulated in the B(disease) group (∆PTGR > 0), and representing the increased RNA stability in the disease group. Positive and negative ∆PTGR value corresponds to increased RNA stability and reduced RNA stability, respectively.

**Figure 1.**
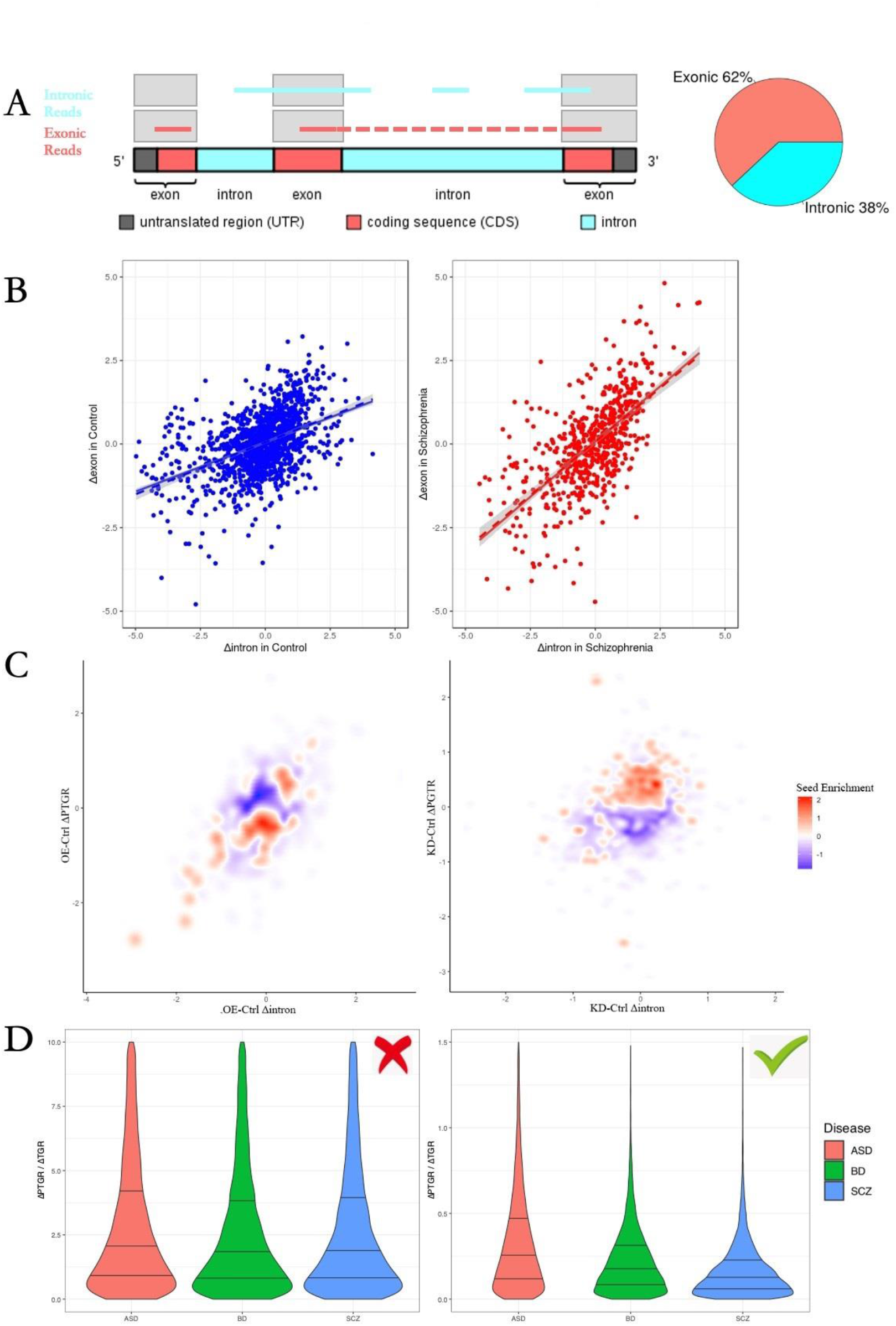
New algorithm for PTGR estimation. (**A)** Theoretical basis schematic diagram of PTGR estimation algorithm (left). The fraction of exonic reads and intronic reads in PsychENCODE bulk RNA-Seq dataset is shows in the pie chart. (right). The intronic reads represent the expression of pre-mRNAs, and the exonic reads represent the expression of mature-mRNAs. **(B)** Schematic diagrams of PTGR estimation method. Scatter plots comparing PTGR of disease (left) and control(right). Y axis represents the changes of exonic reads. X axis represents the changes of intronic reads. Each dot represents a sample, and the solid line represents fitted line. **(C)** Density plot for miR-9 seed sequence enrichments in genes of miR-9 knockdown (left) and miR-9 overexpression (right). Y axis represents the changes of exonic reads. X axis represents the changes of intronic reads. Each dot represents a gene, and the color represents the degree of enrichment of genes with miR-9 seed sequence in the 3’UTR, with red representing enrichment and blue representing depletion. **(D)** Comparison of REMBRANDTS (left) and new PTGR algorithm (right). The violin plot shows the ratio of change of PTGR in disease vs. control to change of TGR in disease vs. control. The three lines within the violin plot represent the 75%, median, and 25% percentiles from up to bottom. ASD, BD and SCZ are respectively displayed in red, green and blue.

To evaluate the performance of our PTGR algorithm, we measured ΔPTGR using RNA-seq data of schizophrenia patient-derived neural progenitor cells with purely post-transcriptional perturbations mediated by microRNAs(*miR-9*) from ref^22^. The regulation of a particular miRNA on its target messages by binding to the miRNA seed region, and therefore the direct targets 3′UTR of miRNAs are enriched in the miRNA seed region. As expected, our results indeed showed genes that were downregulated at the PTGR level in the *miR-9* overexpression condition were significantly enriched for the *miR-9* direct targets, and genes that were upregulated at the PTGR level in the *miR-9* knockdown condition were significantly enriched for the *miR-9* direct targets (Fig. 1C). Our analysis highlighted that the PTGR algorithm is an accurate method for PTGR estimation.

To deeper assess the accuracy of the PTGR algorithm in big data and compared the accuracy relative to other methods, we next applied the PTGR algorithm to the 2160 brain transcriptome samples across three common psychiatric disorders data from the PsychENCODE Consortium. We used the previous algorithm (REMBRANDTS) and the PTGR algorithm to evaluate the influence of PTGR on changes of gene expression (see Methods section). PTGR algorithm revealed the average effect of abnormal PTGR on changes in gene expression levels was approximately 30% of that of abnormal TGR, but the average influence of PTGR estimated by REMBRANDTS on gene expression was 2 times that of TGR (Fig. 1D). The estimation of REMBRANDTS was contradictory to the theory that the majority of the changes in gene expression are driven by transcriptional regulation. Our results demonstrated that the PTGR algorithm should provide more accurate estimates of mRNA stability compared to REMBRANDTS.

### Dysregulation of PTGR in psychiatric disorders

To generate a PTGR architecture across three diseases, we used the PTGR algorithm to assess differential PTGR in ASD, SCZ, and BD compared with control. Intriguingly, we noticed PTGR with the largest changes in ASD compared with SCZ/BD (Fig. 1D), and mRNAs appeared to be significantly more stable in ASD; the other two diseases mRNAs stability showed the reverse pattern (Fig. 2A). We identified widespread differential post-transcriptional regulation genes (DPRGs) in ASD, SCZ, and BD [n =1807, 1019, and 979 genes at false discovery rate (FDR) < 0.05, respectively] (Fig. 2B), indicating a prominent role of post-transcriptional programs in disease development. Besides, DPRGs were enriched for up-regulated PTGR genes in ASD, whereas DPRGs were enriched for down-regulated PTGR genes in SCZ as well as BD. Notably, at the ∆PTGR level, there was inconsiderable cross-disorder sharing of this DPRGs signal, suggesting that ∆PTGR confers a substantial portion of disease specificity. Comparison of ∆PTGR of whole genome genes across three diseases suggested there was a significant SCZ/BD cross-disorder correlation in PTGR changes (Fig. 2B), although the majority of DPRGs were disorder-specific in SCZ/BD. This finding implied that SCZ and BD share overlapping genetic etiology at the PTGR level, and ASD is likely to be with disease specificity at the PTGR level (Fig. 2C). To gain better insights into the contributions of abnormal PTGR in disease development, we subsequently performed Gene Ontology (GO) analysis of ASD-, SCZ-, and BD-associated DPRGs, respectively. Regulation of apoptotic process was enriched in ASD up-regulated DPRGs, implying the involvement of apoptotic in the development of ASD (Fig. 2D). SCZ-/BD up-regulated DPRGs showed significant enriched in immune activation related biological processes (Fig. 2D). Moreover, down-regulated DPRGs were associated with neural development, neuron differentiation, and Synaptic Transmission in SCZ/BD, but no enrichment biological process in ASD (Fig. 2E). Together, this provides direct evidence that the disease specificity and complexity across three disorders partially attributed to the diversity of disease associated mRNA stability, suggesting clear divergence in the pathological mechanisms of psychiatric disorders.

**Figure 2.**
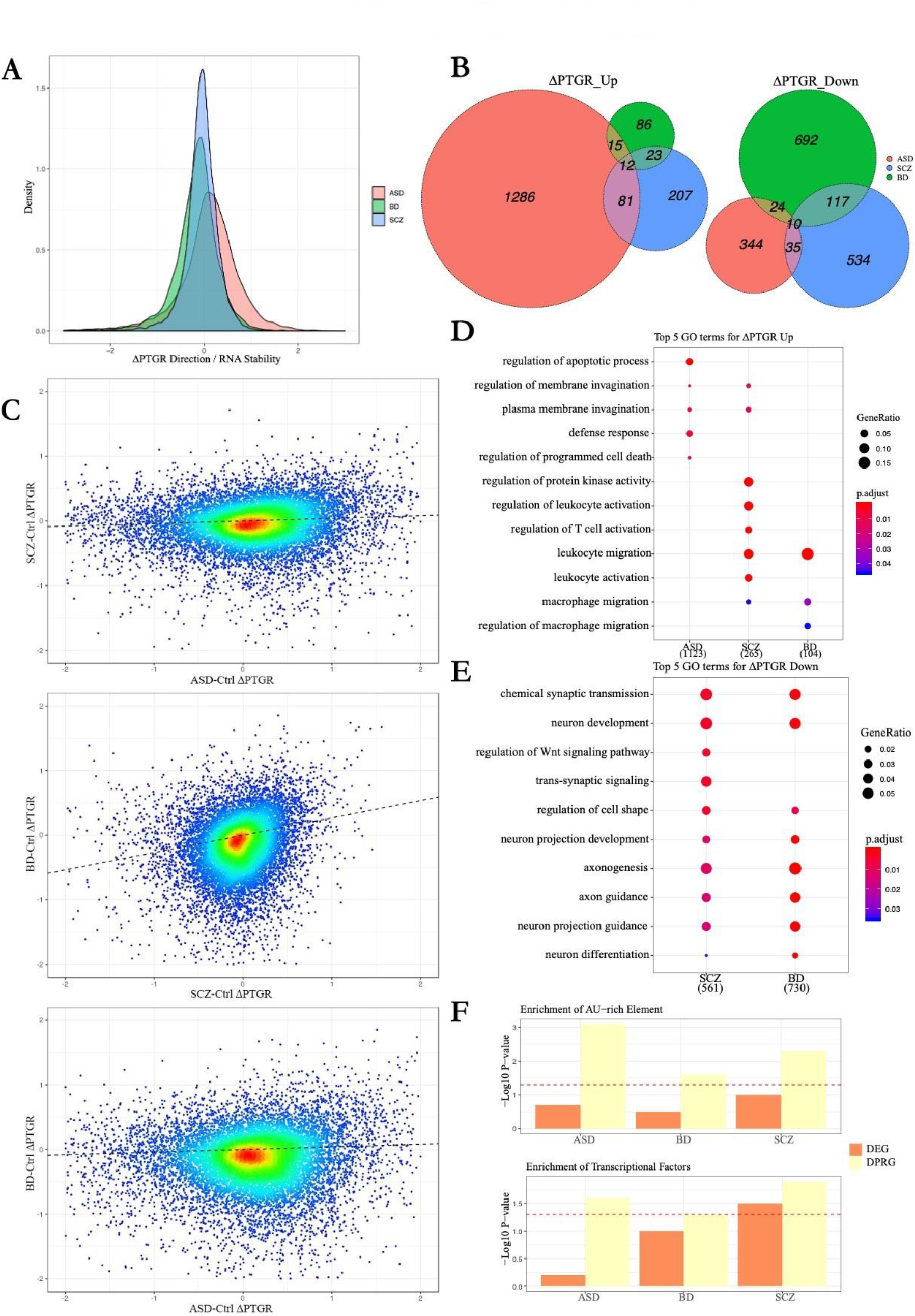
Dysregulation of PTGR in psychiatric disorders. **(A)** Density plot of ∆PTGR distribution within the whole genome genes in three diseases. The ∆PTGR > 0 represents the increase of RNA stability (disease vs. control), whereas ∆PTGR < 0 represents the decrease of RNA stability (disease vs. control). **(B)** Venn diagram showing cross-disorder overlap for genes with significantly differential PTGR (FDR < 0.05). ASD, BD and SCZ are displayed in red, green and blue, respectively. **(C)** The scatter plot shows the degree of correlation of ∆PTGR between diseases. Each dot represents a gene. The color represents the localized density of genes. Red represents high gene density, and the blue represents low gene density. The dotted line represents fitted line. **(D)** Gene ontology enrichments are shown for up-regulated gene of PTGR in each disorder. Numbers in parentheses represent the count of genes in each disorder. **(E)** Gene ontology enrichments are shown for down regulated gene of PTGR in each disorder. Numbers in parentheses represent the count of genes in each disorder. **(F)** The bar plot shows the enrichment degree of regulatory elements AREs(up) and transcription factors(bottom) in DPRGs and DEGs in three diseases.

### Disease-specific key RBPs that modulate mRNA stability in psychiatric disorders

Since RBPs modulate mRNA stability via the Adenylate-uridylate (AU)-rich elements (AREs) and regulate the expression of many genes at the post-transcriptional level^28^, we examined whether DPRGs and differential expressed genes (DEGs) enriched in AREs. We indeed observed ARE enrichment in DPRGs, but not in DEGs (Fig. 2F). This result indicating that RBPs-regulated genes are enriched in DPRGs, and also provides further evidence to support the accuracy of the PTGR algorithm. Additionally, we performed enrichment analysis for transcription factors in DPRGs and DEGs, and observed transcription factors were significantly overrepresented in DPRGs (Fig. 2F), compared with DEGs, suggesting the abnormality of PTGR has a pervasive influence on the whole genome. Although the effect of PTGR on gene expression changes is less than that of TGR, PTGR may contribute as much to the changes of gene expression as TGR by affecting more upstream regulators. This work further highlights PTGR dysregulation as a critical, and relatively underexplored, mechanism linking abnormal gene expression with psychiatric disease pathophysiology.

The above observations prompted us to take a closer look at abnormal mRNAs stability of psychiatric disorders, but it is not clear which factors contribute the most to the abnormal landscape of mRNAs stability of psychiatric disorders. We combined the stability profiles of mRNAs of psychiatric disorders with the binding site predictions of the RBPs, and used multiple linear regression to identify key RBPs that modulate mRNAs stability in psychiatric disorders (see Methods for more details). This analysis identified multiple RBPs that were significantly predictive of disease mRNAs stability (Fig. 3A). Notable examples include FMR1^29^ and QKI^30^, which is known ASD risk gene, and SCZ risk gene^29,31^, respectively. These promising candidate RBPs also including neurodevelopment associated ELAVL3^32^ and IGF2BP family^33^, neuroimmune related gene ZPF36^34^, classical splicing factor SRSF1^35^, and so on. Besides, we observed ELAVL3 was upregulated in ASD, ELAVL3 targets were more likely to be bound and more stable in ASD, and ELAVL3 was downregulated in SCZ/BD, ELAVL3 targets were more likely to be bound and less stable in SCZ/BD (Fig. 3A). Therefore, the presence of 3′ UTR binding sites for ELAVL3 was significantly associated with increased mRNAs stability in three disorders. In sum, the direction of the changes in stability was consistent with the direction verified experimentally in the literature^30^.

**Figure 3.**
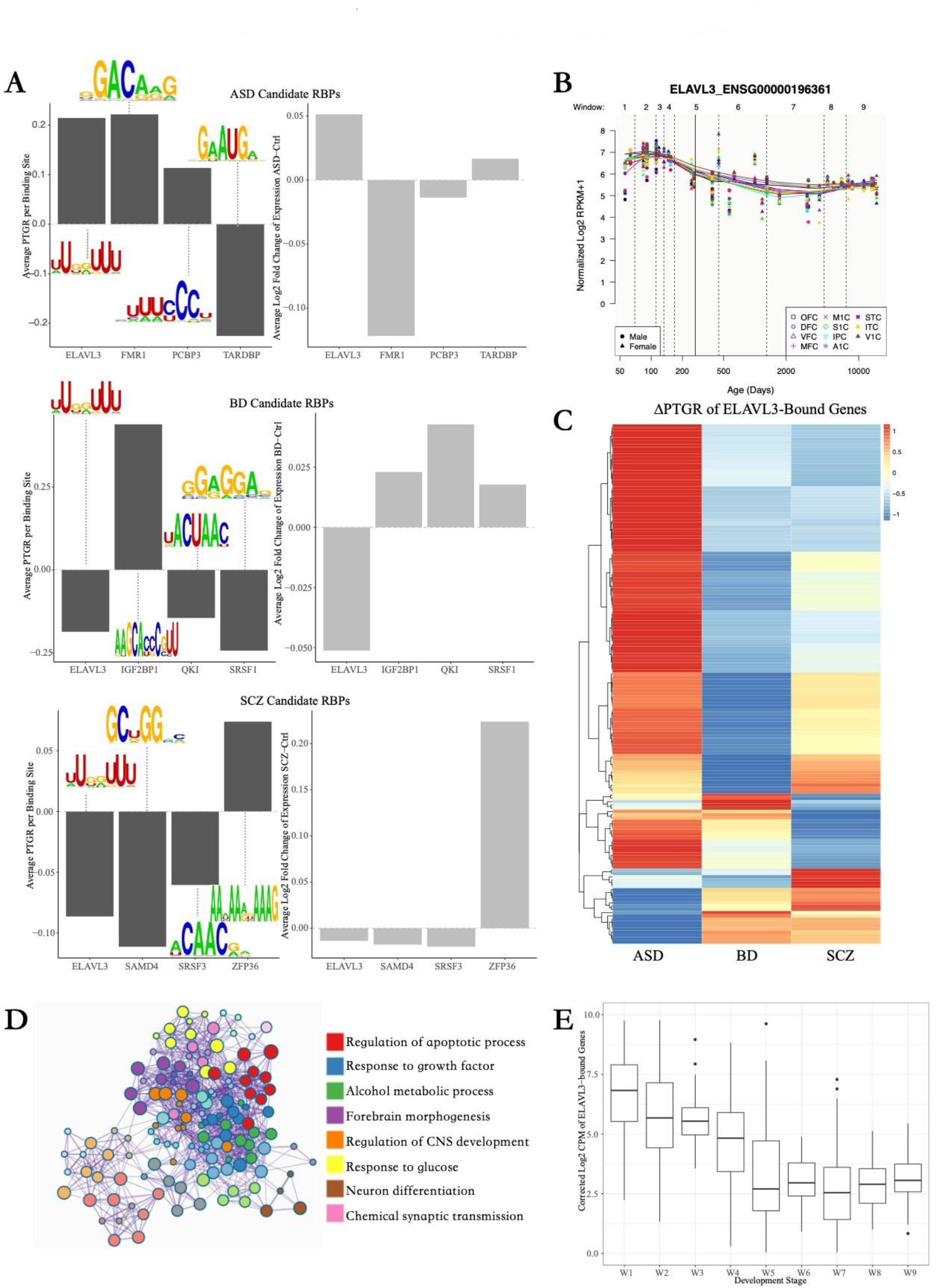
Disease-specific top candidate RBPs that modulate RNA stability in psychiatric disorders. **(A)** RBPs that are significantly associated with disease-specific mRNA stability are shown (FDR < 0.05, *t-test* of regression coefficients). From up to bottom, there are identified RBPs in ASD, BD and SCZ. The left bar plot shows the predicted key RBPs, the corresponding RBP binding motif. Y axis within the left bar plot represents the average PTGR of predicted RBPs-bound genes in disease vs. control. The right bar chart shows the average log2fold-change of the expression of RBP in disease vs. control. **(B)** Plot of spatiotemporal expression of ELAVL3 in human brain. **(C)** Heatmap shows the relative RNA stability of 159 predicted ELAVL3-bound genes in three diseases. The ∆PTGR represents the changes of PTGR of 159 predicted ELAVL3-bound genes in disease vs. control. **(D)** Enrichment analysis of ELAVL3-bound genes. Each color represents a kind of biological process category. **(E)** The box plot shows the average expression of 159 ELAVL3-bound genes at different developmental stages. The stages windows of W1-9 correspond to the developmental time windows in Figure 3 (B).

Intriguingly, ELAVL3, a neural-specific RNA-binding protein which expression centered on early brain development (Figure 3B), displayed alterations of expression in all three psychiatric disorders. We subsequently analyzed 159 ELAVL3-bound genes expression level and found that the average expression of ELAVL3-bound genes also high in brain development (Figure 3E), demonstrating that the involvement of PTGR of ELAVL3 in early brain development. we further examined the ∆PTGR of 159 ELAVL3-bound mRNAs and observed that these mRNAs were mainly increased stability in ASD and reduced stability in SCZ/BD (Fig. 3C). The GO analysis of the ELAVL3-bound mRNAs showed that these genes were significantly enriched in the biological processes of neurodevelopment, regulation of apoptotic process, neuron differentiation (Fig. 3D). Overall, our study implied the disruption of ELAVL*3* may be one of the predominant risk factors that contribute to psychiatry-specific abnormal PTGR and provides a resource for mechanistic insight and therapeutic development.

## Discussion

In this study, we present a new algorithm and integrated it into the transcriptome big data of the brain across three major psychiatric disorders to reveal the abnormal pattern and significance of PTGR for the first time.

The PTGR algorithm can be performed with any type of RNA-seq data set, for example, polyA RNA-Seq, single-cell RNA-Seq, whole RNA-Seq, to gain insights into the post-transcriptional regulatory mechanism responsible for the changes of gene expression. Notable, the accuracy of the PTGR estimate depends on the larger sample size, our method is more suitable for large sample data.

RNA stability profiles across three diseases displayed that the aberrant PTGR changes give a new characteristic of each disorder, and PTGR changes have expanded the understanding of the pathogenesis of psychiatric disorders. The similar pattern of RNA stability between SCZ and BD in disease revealed their convergence at the PTGR level, and the reverse pattern on the RNA stability of between ASD and SCZ/BD showed the divergence of PTGR across three disorders. According to the shared and specific PTGR abnormalities of three diseases, we believe that SCZ and BD share the pathogenic mechanism, and the pathogenic mechanism of ASD is specific more than SCZ and BD. The convergence and divergence of mRNA stability across three diseases provide the theoretical basis for investigations targeting shared and specific disease mechanisms to link causal drivers with psychiatric disorders.

Gene expression and alternative splicing events are determined by both TGR and PTGR. However, there are limited studies to accurately assess the contributions of TGR and PTGR on changes in gene expression in psychiatric disorders. Our results found the effect of PTGR abnormality on gene expression in three common psychiatric disorders was 30% of that of TGR abnormality, suggesting that TGR changes exhibit larger effect sizes in diseased brain gene expression than TGR changes. However, our results found that the upstream regulatory factors were enriched in the DPRG, suggesting that abnormal PTGR in psychiatric disorders leads to abnormal expression of the upstream regulatory factors, which indirectly has a significant impact on downstream gene expression. Therefore, we hypothesized that PTGR and TGR might contribute equally to changes in gene expression of psychiatric disorders, and this finding brought the significance of the PTGR in psychiatric disorders to a higher level.

The current understanding of the relationship between psychiatric disorders and PTGR mainly comes from the research of microRNAs (miRNAs), which regulate the degradation and translation of hundreds of mRNAs by binding to the 3 ‘non-coding region of mRNAs. Numerous well-characterized miRNAs have been associated with psychiatric disorders, including *miR-132, miR-195, miR-18a, miR-137*, and so on. Although the influence of miRNAs and RBPs on PTGR has not been determined, existing studies have shown that RBPs are the all-around regulator of mRNA metabolism, and these findings suggest that RBPs play a non-negligible role in PTGR. Moreover, compared with transcription factors, RBPs regulate the end products of expression more directly, making RBPs a “new favorite” for drug intervention targets. The human genome encodes more than 800 RBPs involved in the PTGR of mRNAs, yet only a small fraction has been functionally characterized. Encouragingly, our results identified many promising potential psychiatric susceptibility RBPs at the post-transcriptional regulation level. *Ince-Dunn* et al. performed a genome-wide analysis of *nElavl* targets and revealed that one function of *nElavl* is to control excitation-inhibition balance in the brain^32^. Our results indicated ELAVL3 is a candidate RBP that plays a critical role in three psychiatric disorders, its activity increased in ASD, and in SCZ/BD activity decreased, resulting in cell apoptosis, neural development, and neuron differentiation of related genes mRNA stability aberrance. These observed suggesting that the PTGR mediated by ELAVL3 may be related to the excitation-inhibition imbalance and play an important role in the development of psychiatric disorders. Our results can help clarify the regulatory mechanism of RBP-mediated PTGR.

In summary, we applied a new PTGR estimation method to RNA sequencing data of 2160 brain samples from ASD, SCZ, and BD, as well as controls individuals. We found that SCZ and BD shared a similar abnormal pattern of the PTGR, whereas ASD has a specific abnormality of PTGR, revealing the regularity of PTGR abnormality in three psychiatric disorders. Besides, our results identified RBPs that play a critical role in psychiatric disorders, suggesting that the abnormalities of RBP-mediated PTGR may contribute to the development of psychiatric disorders. These findings bring new strategies for mechanistic insight and disease therapeutic development.

## Author contributions

The study was designed by M.L.. M.L. and L.L. collected data. M.L., L.L. and

Y.W. did the bioinformatic analyses. Y.W., M.L., and L.L. wrote the manuscript.

## Acknowledgements

We are grateful to all laboratory members for comments on the manuscript.

